# Phase-targeting rapid cryofixation of the beating heart and histological analysis unveil contractile state-dependent sarcomere dynamics

**DOI:** 10.1101/2025.11.15.688585

**Authors:** Shoko Tamura, Kentaro Mochizuki, Yasuaki Kumamoto, Yuma Morishita, Masahito Yamanaka, Wen Jin Ho, Yoshinori Harada, Katsumasa Fujita, Hideo Tanaka

## Abstract

The heart is a functional syncytium consisting of numerous cardiomyocytes that repetitively exhibit coordinated contractions/relaxations. However, the extent to which myocyte sarcomere arrangements in the heart differ across beats is unknown. To examine this, we conducted cardiac phase-targeting rapid cryofixation of Langendorff-perfused rat hearts. We adjusted the timepoint of cryogen exposure to the electrically paced heart and observed phase-dependent differences in the sarcomere length (SL) of subepicardial myocytes by α-actinin immunohistochemistry, namely a significantly shorter SL during systole than during diastole. We detected spatially inhomogeneous SL distributions by generating a heatmap of the myocardium. For peak systole the SL heatmap exhibited nearly uniform SL shortening within and among the individual myocytes with some myocardia exhibiting nonuniform SLs. During diastole, the heart showed predominant SL elongation, which was also accompanied by patchy distributions of locally short-SL regions, reflecting inhomogeneous SLs. This SL inhomogeneity was attenuated by pharmacological relaxation by 2,3-butanedione monoxime. The heatmap of the rapidly-frozen heart during ventricular fibrillation also revealed inhomogeneous SLs within and among individual myocytes. Overall, cardiac phase-targeting cryofixation unveiled in-depth behaviors on SL in the heart. Our cryofixation strategy will open a new horizon to clarify precise spatiotemporal changes in sarcomere structures and understand cardiac functions.

## Introduction

The heart is a functional syncytium consisting of numerous cardiomyocytes that repetitively exhibit coordinated contraction and relaxation [**1–3**]. However, the extent to which the sarcomere structures of individual myocytes change in the working heart on a beat-by-beat basis is unknown. Although chemical fixation using formalin-based fixatives e.g., paraformaldehyde (PFA) [**4–6**], has been widely used to visualize myocyte structures, no information is available on the precise cardiac phase-dependent histology of myocyte sarcomere arrangements within the heart because of the inability to determine the timepoint of fixation in the cardiac phase, e.g., during systole or diastole. For more than three decades technological advancements in fast fluorescence, live-cell imaging have enabled us to understand the spatiotemporal structures and/or functions of beating hearts, e.g., intracellular Ca^2+^ dynamics and cell shape [**7–12**].

However, obtaining convincing cardiac phase-specific histology of the myocytes in the beating heart with an adequate signal-to-noise ratio is still difficult, because of the trade-off between the temporal resolution and signal strength, which affects the spatial resolution and field of view during measurement. Fukuda’s group successfully visualized rapid structural changes in sarcomeres in beating hearts *in situ* under fluorescence microscopy by targeting a few myocytes [**12**]. For histological assessment, however, simultaneous detection of larger number of myocytes would be required.

We assumed that if the heart could be “instantaneously fixed” in an arbitrary cardiac phase by rapidly freezing the heart, convincing histological information would be obtained on an extensive area of phase-specific sarcomere structures in the myocardium. The rapid-freezing technique, which was originally established for electron microscopy to fix the microstructures of living samples [**13–16**], has recently been applied to the field of optical microscopy to achieve a high signal-to-noise ratio with high spatial resolution [**17–20**]. This technique has also been advanced as a method for the instantaneous fixation of moving samples in a targeting phase to preserve its structure and function, e.g., eelworm, cancer cell lines, isolated cardiomyocytes, for multimodal observations before and after rapid freezing in the targeted phase [**17–20**]. In particular, we have recently succeeded in instantaneously fixing the cellular structures and intracellular Ca^2+^ waves in isolated neonatal rat isolated cardiomyocytes at specific time points with a temporal precision of ±10 ms during optical observation using a rapid-freezing technique [**20**].

In this study, we aimed to develop a rapid-freezing system to fix the perfused rat heart during arbitrary cardiac phases, particularly during systole and diastole, to understand the precise phase-dependent sarcomere structures of individual cardiomyocytes in the heart. Rapid freezing was also applied to the heart during ventricular fibrillation.

## Results

### Cardiac phase-targeting rapid-freezing system

Figure 1A shows a schematic illustration of the phase-targeting rapid-freezing system for the Langendorff-perfused rat heart. The system (Fig. 1B, top left panel) consists of two parts: a cryogen-ejection component (upper part) and a sample stage component with a heart chamber (lower part). The cryogen-ejection system consists of a tank to store liquid propane as a cryogen and a TTL trigger-controlled electromagnetic valve for opening and ejecting the cryogen. On the bottom of the sample chamber the perfused heart was placed with the left ventricular (LV) surface upward under consecutive pacing from the ventricular apex at 0.5 Hz. A ring-shaped aluminum electrode was gently placed on the LV surface for ECG recording and for the detection of the cryogen reaching the heart (Fig. 1B, bottom panel). We cryofixed the heart during peak systole or end diastole by adjusting the time point of valve opening for cryogen ejection (Fig. 1C; for details *see the “Methods”*).

**Figure 1.**
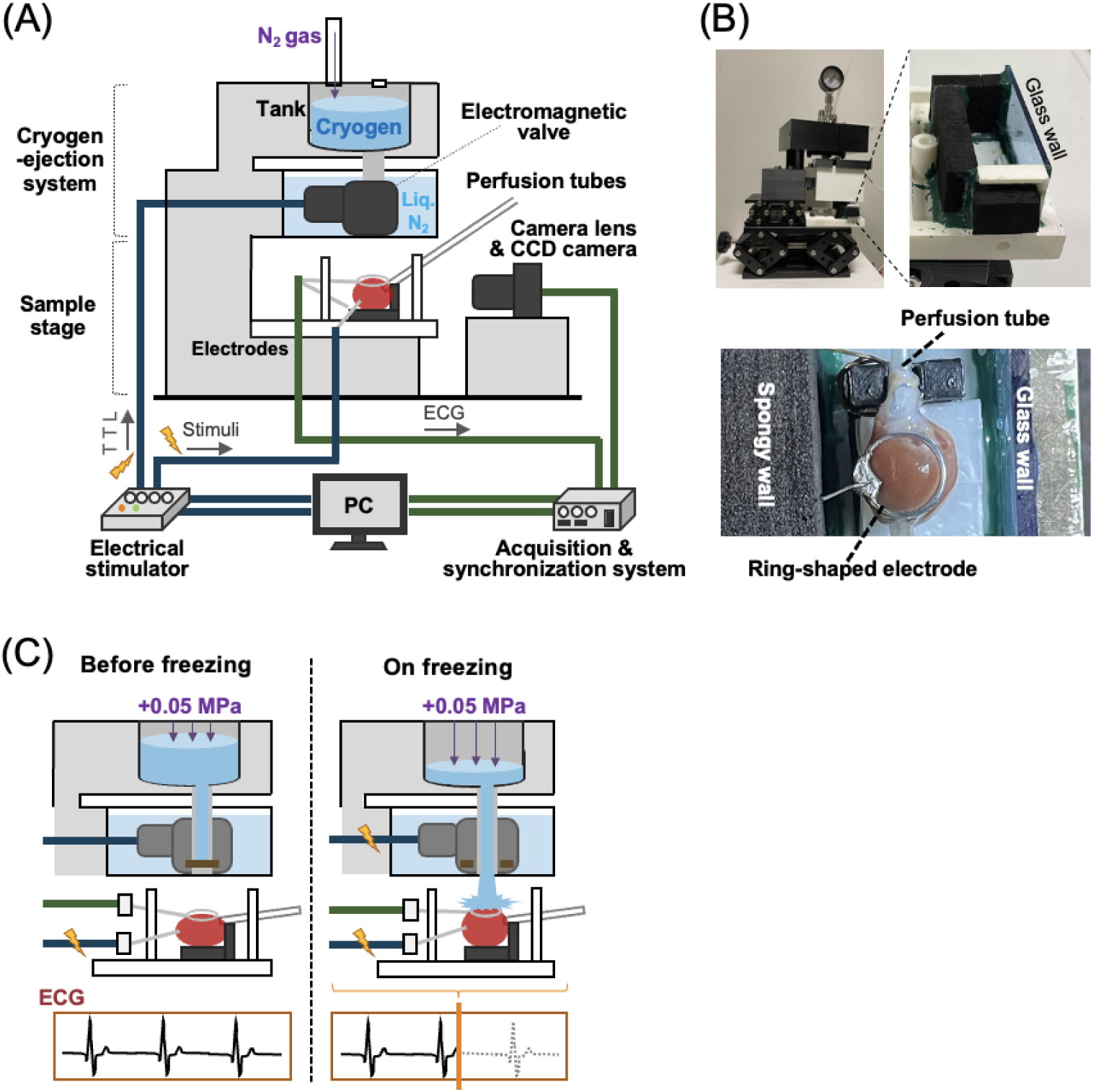
Diagram of the rapid-freezing system for the perfused rat heart. (A) Overall view of the system. (B) Photos showing the main body in the system composed of the cryogen-ejection system and sample stage (upper left panel), the sample chamber on the sample stage (upper right panel), and a top-down view of the heart situated in the chamber with the ring-shaped electrode on the left ventricular surface (bottom panel). (C) Schematic illustrations of the system before (left panel) and during (right panel) the rapid freezing of the heart. The gauge pressure of 0.05 MPa was applied into the cryogen tank. ECG: electrocardiogram.

### Phase-targeting rapid freezing of the heart

Representative side-view images of the hearts in the process of rapid freezing during peak systole (**A**) and end diastole (**B**) are shown in Fig. 2. While heart contraction was recorded from the side with a high-speed CCD camera at 143 frames/s, the heart was instantaneously exposed to the cryogen during peak systole. Sequential images show that the heart surface was rapidly exposed to the cryogen (Fig. 2A**-i**; *also see* **Supplementary Video 1** online). The image showing the vertical shift image of the ring electrode on the top of the heart over time clearly indicates sequential changes in contractile motion (Fig. 2A**-ii**). Evidently, the heart was instantaneously exposed to the cryogen during peak systole, as evidenced by an abrupt disappearance of the heart surface (shown by an arrowhead) with a large deflection of the ECG trace upon cryogen ejection (Fig. 2A**-iii**). Rapid freezing of the heart was also conducted during end diastole, i.e., with targeting at 1950 ms after stimulation (Fig. 2B). Side-views and sequential changes in the heart surface are presented for all the hearts cryofixed during peak systole and end diastole in **Supplementary** Fig. 1.

**Figure 2.**
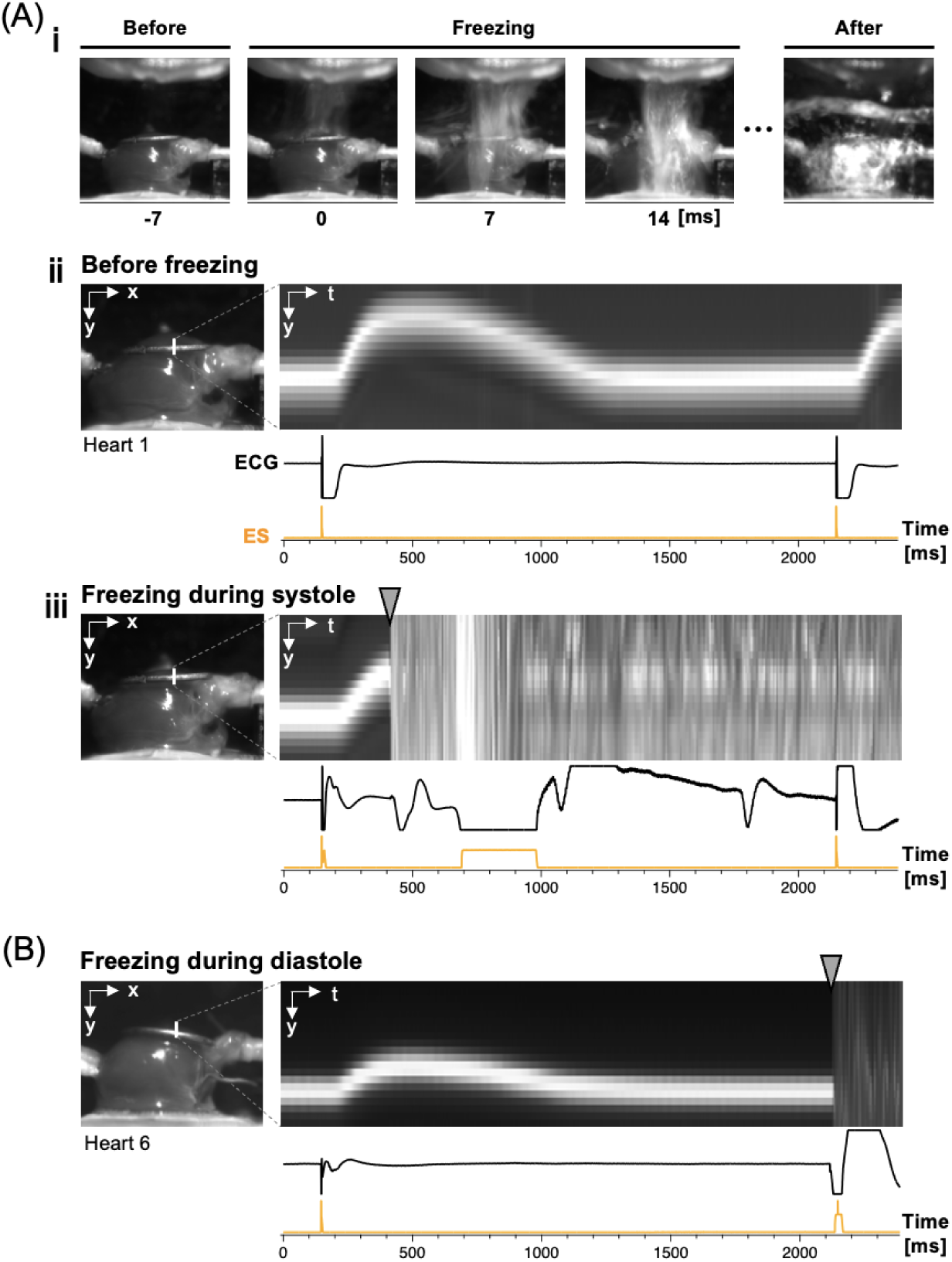
Side views of the heart during rapid freezing. (A-i) Sequential images captured 7-ms before, immediately upon (0 ms), during (7 ms and 14 ms), and after the heart was exposed to the cryogen during peak systole. (A-ii) Side view of the heart (left panel: at end diastole) and image showing the vertical shift of the ventricular surface (ring electrode) over time during the cardiac phase (right panel) with the corresponding traces of electrocardiogram (ECG) and electrical stimulation (ES: 0.5Hz) below. (A-iii) Rapid freezing of the heart during peak systole. Side-view image of the heart (left panel: at end diastole) before cryogen exposure and an image of the vertical shift of the ventricular surface with rapid cryogen exposure during peak systole. Note the abrupt disappearance of the ventricular surface with a large deflection of the ECG trace upon cryogen exposure (arrowhead). (B) Rapid freezing of the heart during end diastole. Side-view image of the heart (left panel: at end diastole) before cryogen exposure and an image of the vertical shift of the ventricular surface with rapid cryogen exposure at end diastole (arrowhead).

### Histological evaluation of the rapidly-frozen heart

For histological evaluation of the cryofixed heart we performed freeze substitution after rapid freezing: the frozen heart was gradually fixed with PFA and acetone under sequential temperature elevation to room temperature and the cryofixed microstructure maintained [**21, 22**] (for details, *see the* “*Methods”*). Gross observations indicated that the freeze-substituted heart did not obviously differ from the heart immediately after rapid freezing (Fig. 3A). Vertical and horizontal cross sections of the LV surface (Fig. 3B) revealed differences in hematoxylin and eosin (HE)-stained histology depending on the tissue depth and the time point of cryogen exposure (Fig. 3C, D). The rapid-freezing procedure preserved the myocyte structure only up to the limited depth of the ventricular surface. As shown in the images of vertical-sections obtained from underneath the cryogen-exposed surface and the corresponding binarized images (Fig. 3C, top panels), a limited number of small vacuoles (< 1 µm) were observed in the myocytes located at an up to approximately 50 µm depth from the surface, whereas vacuoles were entirely distributed in myocytes located deeper than approx. 50 µm, probably due to ice-crystal formation [**23**]. With respect to the myocytes beneath the ring electrode (i.e., without direct cryogen exposure), more and larger vacuoles were observed on the outermost surface (Fig. 3C, bottom panel).The myocyte structure was preserved on the horizontal planar surface of the direct cryogen-exposed heart, as the subepicardial myocardium showed no clear vacuoles or tissue deformation of the heart that was rapidly frozen (RF) during peak systole (hereafter, “RF-systole”) or end diastole (“RF-diastole”) (Fig. 3D). However, no clear difference was observed between the images of HE-stained “RF-systole” and “RF-diastole” hearts or the sole PFA-fixed heart (“PFA”, Fig.3D, bottom panel) because of the lack of visible striations in the myocytes. For an additional comparison, myocytes showed marked granular patterns in the rapidly frozen heart once it was thawed at room temperature and subsequently underwent rapid refreezing (**Supplementary** Fig. 2A). Such abnormal patterns are caused by ice-crystal formation in the process of relatively slow temperature changes during the process of thawing [**23**]. Subcellular vacuoles were also remarkable in the fluorescence images of the heart that was rapidly frozen again after thawing (**Supplementary** Fig. 2B, right panel, F-actin staining). In addition, the fluorescence images for Z-lines (α-actinin) and F-actin clearly revealed the deformation of the cell structure of the RF heart after thawing [**24**] compared with the RF heart without thawing (for comparison, *see* Fig. 4). Thus, due to the very low degree of vacuolar formation and indiscernible cellular deformation in a limited area of the surface of the ventricle, rapid freezing of the beating heart would be possible for histological evaluation at least on the spatial scale of optical microscopy. However, images of HE-stained sections of the rapidly-frozen heart revealed no obvious differences in the myocyte structures between the systolic and diastolic phases.

**Figure 3.**
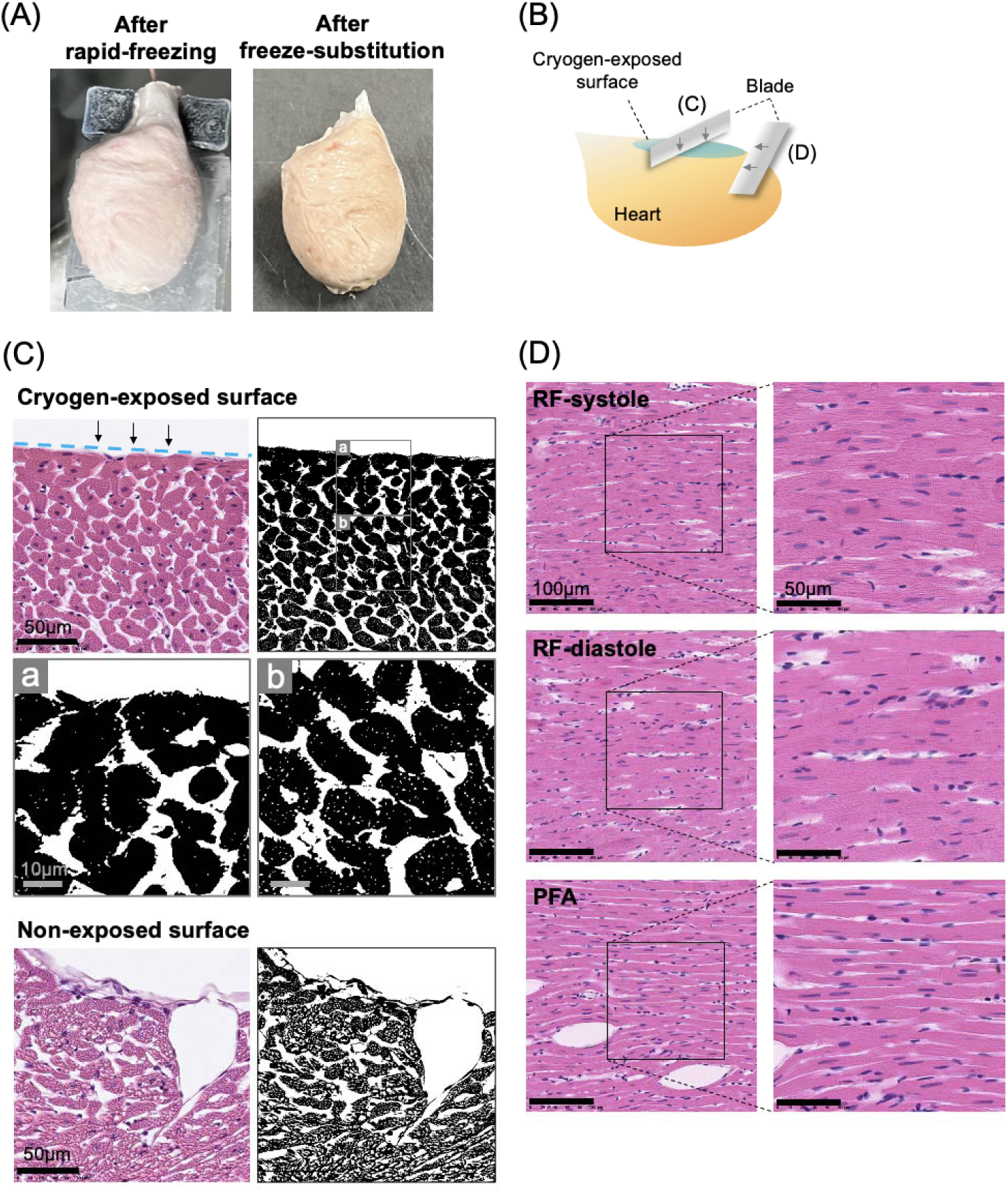
Gross and microscopy images of the rapidly-frozen heart. (A) Gross appearance of the rapidly-frozen hearts, immediately after rapid-freezing (left panel) and after subsequent freeze-substitution (right panel). (B) Schematic illustration showing directions of slicing of thin sections for HE-staining. (C) Cross-sections underneath the cryogen-exposed surface (inside the ring of the ring-shaped electrode on the left ventricle) and nonexposed surface (underneath the ring-shaped electrode). The corresponding binarized images are shown to emphasize the presence of small vacuoles. For the cryogen-exposed surface, enlarged images of the binarized image are shown up to an approximately 50 µm depth from the surface (a) and that depth is greater than approximately 50 µm (b). (D) The en face plane near the cryogen-exposed ventricle surface (<30 µm) of the heart during peak systole and end diastole and the nonrapidly-frozen heart fixed by PFA perfusion. Thickness of the sections: 4 µm.

**Figure 4.**
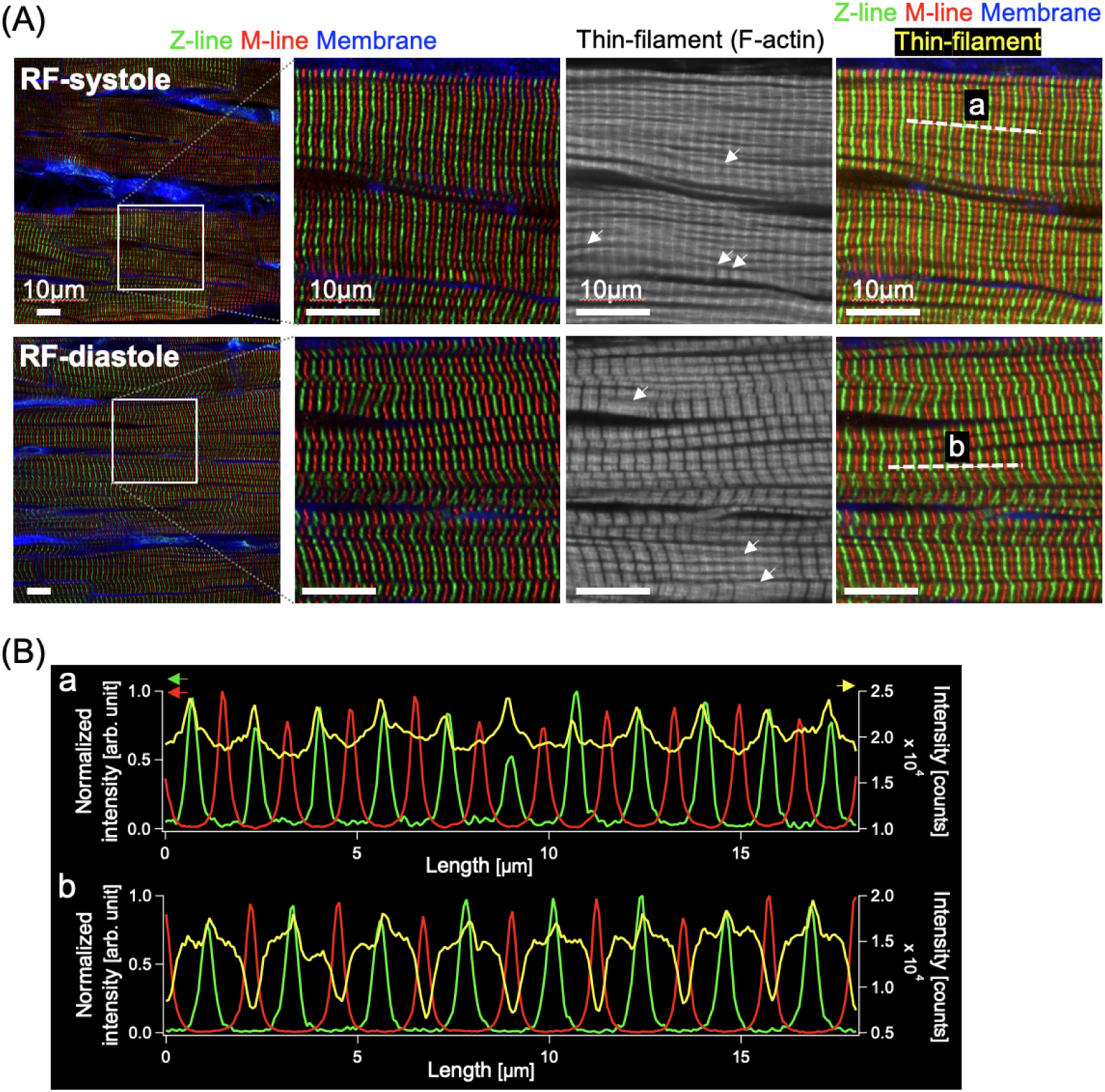
Confocal fluorescence images of the subcellular myocyte structures of the cryofixed hearts that were rapidly frozen during peak systole (“RF-systole”) and end diastole (“RF-diastole”). (A) Fluorescence images of the Z-lines, M-lines, thin filaments (F-actin), and cell membranes labeled with immunofluorescence staining for α-actinin and titin, and fluorescence labeling with phalloidin and wheat germ agglutinin (WGA), respectively. The images in the 2^nd^ to 4^th^ columns from the left are cropped and enlarged from the white square region in the leftmost images. (B) Averaged intensity line-profiles of the Z-line (green), M-line (red), and thin filament (yellow) extracted along with the white-dotted lines (a) and (b) in the images in the fourth column of (A). The profiles were averaged from 3 adjacent line profiles.

### Different sarcomere structures between systole and diastole

We next conducted fluorescence imaging of the subepicardial myocardium of the rapidly-frozen heart to assess the cardiac phase-dependent differences in the myocyte structure. The Z-lines and M-lines of the sarcomere were subjected to immunofluorescence staining for α-actinin and titin, respectively. In addition, the thin filaments (F-actin) and the cell membrane were stained with phalloidin and WGA, respectively (*for details, see in the “Methods”*). Confocal fluorescence images of the subepicardial myocardium (<30 µm from the surface) revealed periodic arrangements of the Z-lines (green) and M-lines (red) in individual myocytes in both the RF-systole and RF-diastole hearts (Fig. 4A). The spatial distribution of the thin filaments shown in grayscale differed markedly between the two cardiac phases: for the RF-systole myocyte, no discernible gaps were identified between the thin filaments, whereas wide gaps were definitively identified for the RF-diastole myocyte. Notably, some small parts of the thin filaments revealed paradoxically opposite behaviors, i.e., wide gaps for RF-systole and no gap for RF-diastole as indicated by white arrows. Figure 4B shows the profiles of the fluorescence intensity distribution for the Z-lines (red), M-lines (green), and thin filaments (yellow) along the longitudinal direction of the myocardial fibers defined as (a) and (b) in Fig. 4A. The locations of the Z- and M-lines were confirmed as peaks, whose numbers were approximately 11 and 10 for RF-systole and 8 and 7-8 for RF-diastole, respectively. Consistent with the fluorescence images of the thin filaments (yellow), the line profiles for the RF-diastole myocyte showed deep cleavage furrows corresponding to the gaps, whereas those for the RF-systole myocyte showed no discernible cleavage furrows (Fig. 4B). In addition to the Z-lines, M-lines, and thin filaments, the distribution of thick filaments was also visualized using myosin (heavy chain) immunohistochemistry (**Supplementary** Fig. 3); as shown in the thick filaments (cyan) and thin filaments (magenta) the locations of the A-, I-, and H-bands were distinguished in the RF myocardium. Overall, rapid-freezing fixation of the heart was successfully accomplished during systole and diastole, and fluorescence images of the rapidly frozen heart allows us to distinguish sarcomere structures between RF-systole and RF-diastole heart tissues.

### Quantification of the sarcomere length (SL) in rapidly frozen hearts

We quantitatively analyzed the SL, the basic ultrastructural unit in the contraction of cardiomyocyte [**25**], in the subepicardial myocardium of the RF hearts during systole and diastole to further clarify the cardiac phase-dependent differences in the myocyte sarcomere configuration (Fig. 5). The Z-lines were identified by immunofluorescence staining for α-actinin (green) along with cell membrane staining with WGA-conjugated fluorescent dyes (blue) and, in some cases, staining of the intercalated disc using N-cadherin histochemistry (red) for the identification of cell borders (Fig. 5A**-i**). The SL was measured as the interval of the neighboring Z-lines of the individual myocyte using fast Fourier transform (FFT) based on a previously reported method [**9**] (*see* details in the *“Methods”*). As shown in Fig. 5A**-ii**, sharp peaks of the intensity profiles for α-actinin fluorescence along the white line (labeled “a”) indicate the spatial locations of the Z-lines. Figure 5A**-iii** shows the superposition of the representative amplitude spectra for SL along the line profiles a-h for RF-systole and a-i for RF-diastole as solid lines and dotted lines, respectively. The peak position of the amplitude spectra on the cycle length axis (µm) was regarded as the interpeak interval (i.e., representative SL) of the corresponding myocyte. Quantitatively, the mean SL values from 20 to 25 different myocytes belonging to the individual hearts were significantly shorter for RF-systole (n = 5 hearts) than for RF-diastole (n = 5 hearts) (Fig. 5B). The mean SL value for the RF-diastole heart was not significantly different from the value for the RF-diastole heart under mechanical relaxation induced by 2,3-butanedione monoxime (BDM) (“RF-diastole (BDM)”; n = 3 hearts) [**26**], and was similar to the value for sole PFA fixation (“PFA”; n = 3 hearts). The means and standard deviations of SL were 1.57 ± 0.12 µm for RF-systole, 1.93 ± 0.14 µm for RF-diastole, 1.97 ± 0.11 µm for RF-diastole (BDM), and 1.82 ± 0.11 µm for PFA. We next plotted all the SLs measured from individual myocytes with different symbols and colors depending on the heart to further explore the distribution of representative SLs in each heart (Fig. 5C). The significance of differences in the data set of the SL plots from individual cells was evaluated using a linear mixed effect (LME) model [**27, 28**] to account for the hierarchical property of the data. The differences in the SLs of individual myocytes were the same as those analyzed on a heart-by-heart basis, as shown in Fig. 5B. Additionally, the mean SL for the rapidly-frozen heart after PFA fixation (“PFA+RF”) (1.77 ± 0.09 µm) appeared slightly shorter than that of the sole PFA-fixed heart (“PFA”) (1.82 ± 0.11 µm) (**Supplementary** Fig. 4); however, no significant difference was observed between the two, indicating that procedures of the rapid freezing and subsequent freeze substitution would not affect the SL. Thus, as expected, rapid-freezing fixation of the LV heart revealed quantitatively shorter myocyte SLs during systole than during diastole.

**Figure 5.**
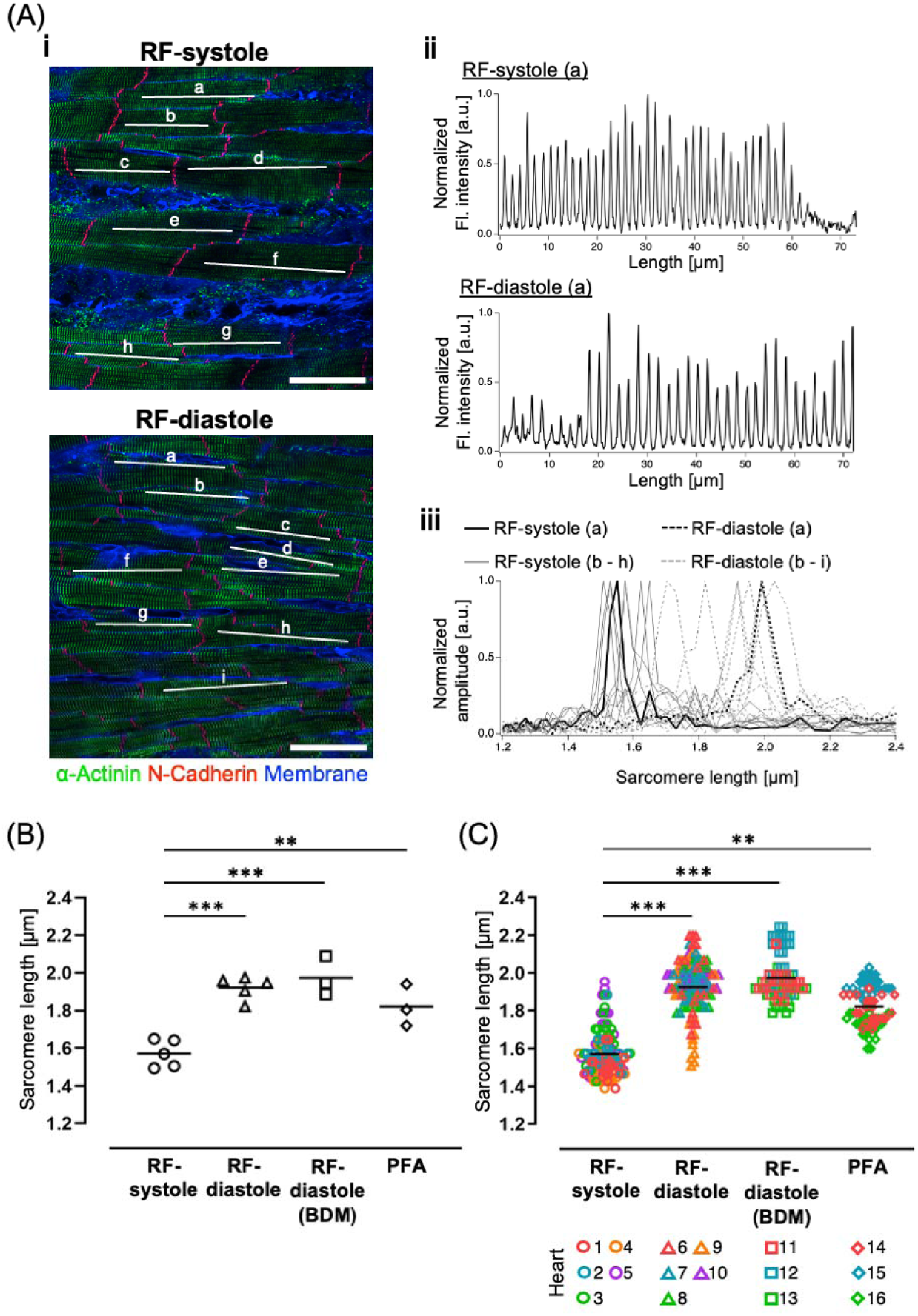
Quantitative results for the myocyte sarcomere length (SL) in the cryofixed heart during systole and diastole based on a spatial frequency analysis using fast Fourier transform (FFT). (A-i) Fluorescence images of α-actinin, N-cadherin, and the cell membrane of rapidly-frozen hearts during peak systole (“RF-systole”) and end diastole (“RF-diastole”), as measured with a confocal laser-scanning microscope. Scale bars: 50 µm. (A-ii) Representative intensity profiles obtained along the lines labeled as (a) in the fluorescence images in A-i. (A-iii) Amplitude spectra of each line profile in the fluorescence images, calculated using FFT. The peak positions are adopted as the representative SL at the sampled myocytes. (B, C) Statistical results for the extracted sarcomere length per heart (B: n = 3-5 hearts) and per myocyte (C: n = 20-25 cells/heart. **p<0.01, ***p<0.001.

### Heatmaps revealed spatial varieties of local sarcomere length

We constructed a heatmap for SLs based on the fluorescence images of Z-lines (α-actinin) to understand the histological aspects of intracellular arrangement of sarcomeres (*see the details in the “Methods”*). Representative heatmaps of the LV surface for “RF-systole”, “RF-diastole”, “RF-diastole (BDM)”, “PFA”, and “PFA+RF” hearts are displayed with color variation depending on the local SL in Fig. 6 (for all analyzed hearts, attached in **Supplementary** Fig. 5). The map was reconstructed by plotting the representative SL value (1.0-2.6 μm) for each pixel as a color grade, which was designated by the FFT for fluorescence intensity distribution on the neighboring 100-pixel width (i.e., 7.0 µm). As shown in the three representative heatmaps of the three different RF-systole hearts (Fig. 6A), almost all the cell areas appeared orange to yellow (the corresponding SLs ranged from approximately 1.4-1.8 µm), indicating that the myocytes were contracted. However, not all the myocytes showed spatially uniform contractions: SLs were regionally diverse within and among the individual myocytes. Specifically, the RF-systole heart contained some green-colored regions corresponding to long SLs of approximately 1.8-2.1 µm. In practice, these longer SL regions were noticeably observed in the maps of hearts 3, 4, and 5 in **Supplementary** Fig. 5A. With respect to the RF-diastole heart, local regions colored yellow to green (approximately 1.6-2.1 µm in SL) were dominant compared with the RF-systole heart, although the almost all the myocytes were presumably being relaxed (Fig. 6B). Interestingly, however, locally orange-colored regions indicating short SLs (approximately 1.4-1.6 µm) were scattered on the maps for RF-diastole hearts, as shown on the leftmost- and middle-maps. Such shorter-SL compartments presented in an orange-to-yellow-color were frequently distributed in the maps of the hearts 6, 7, and 9 in **Supplementary** Fig. 5B, with spatial widths ranging approximately 10-50 µm along the longitudinal axis of the myocytes. With respect to the RF-diastole (BDM) hearts (Fig. 6C) and the solely PFA-fixed hearts (Fig. 6D), the myocardia were entirely dominated by yellow-to-green colors (SL of 1.6–2.1 µm) indicating relaxation, but with a few portions of short SL. In more detail, the case of the “PFA” heart showed more dominant yellow regions (i.e., shorter-SL region, heart 14 and 16 in **Supplementary** Fig. 5D) than the RF-diastole (BDM) heart in **Supplementary** Fig. 5C, indicating a tendency to have an entirely shorter SL in the former conditions than in the latter. Additionally, the heatmaps of the hearts that were cryofixed after PFA fixation (“PFA+RF”) (Fig. 6E and **Supplementary** Fig. 5E) did not differ from those of the heart fixed with only PFA (Fig. 6D and **Supplementary** Fig. 5D), indicating that the PFA-fixed heart was adequately fixed close to relaxed conditions with no additional effect of the rapid freezing and subsequent freeze substitution procedures. Together, the SL heatmaps revealed spatial variability in the sarcomere contractile state in the individual myocytes of the heart during systole and diastole, which was considerably attenuated by forcible relaxation induced by BDM and gradual fixation with PFA.

**Figure 6.**
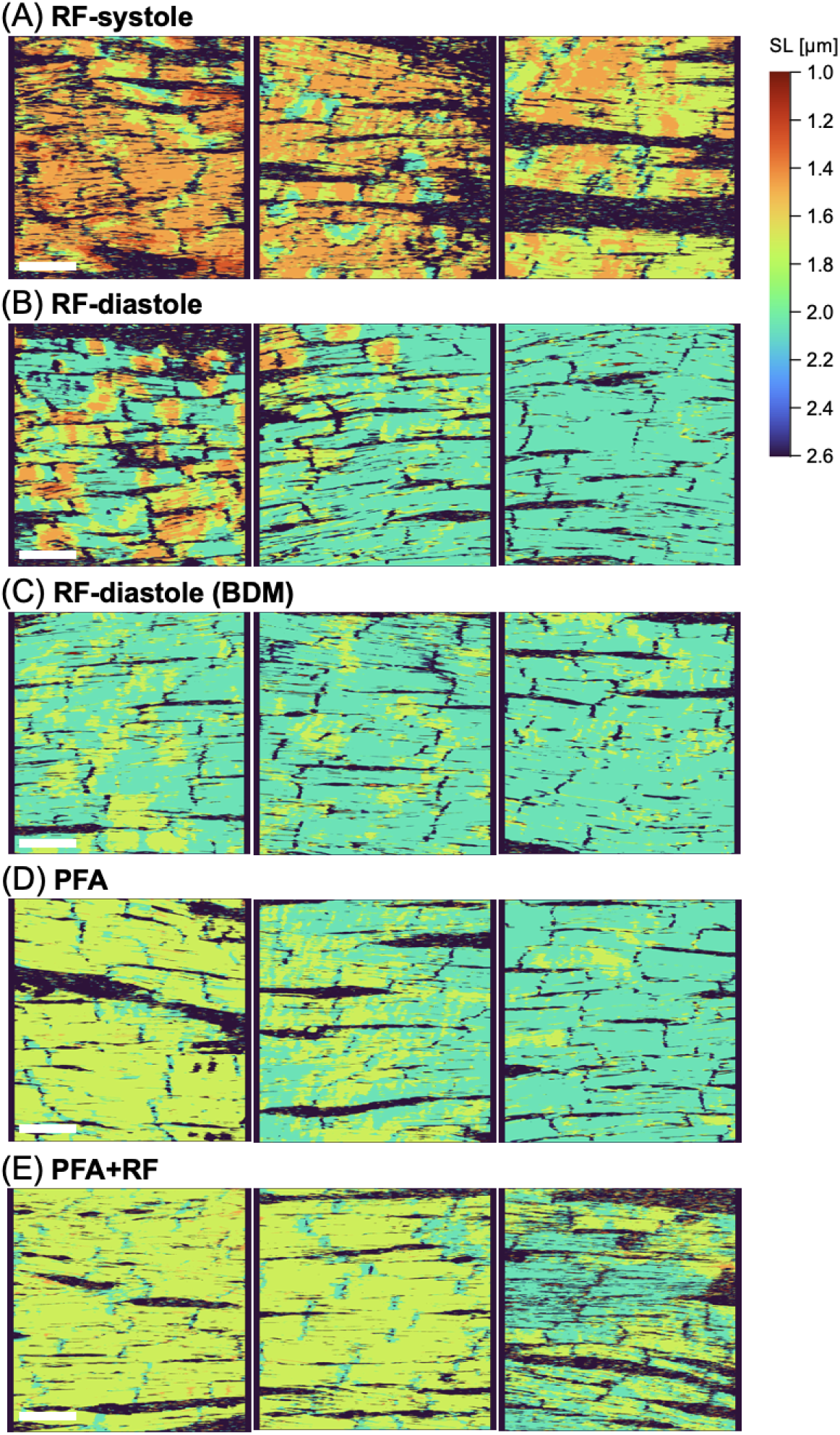
Heatmaps depicting the distributions of local sarcomere lengths (SL), based on the fluorescence images of Z-lines (immunostaining of α-actinin) in the hearts fixed under each condition: rapid-freezing during peak systole (A: “RF-systole”) and end diastole (B: “RF-diastole”), rapid freezing under BDM treatment (C: “RF-diastole (BDM)”), fixation by PFA-perfusion alone (D: “PFA”), and rapid-freezing after PFA fixation (E: “PFA+RF”). Please note that the areas of nonmyocyte or intra-myocyte SL out of the range of 1.0 to 2.6 µm are colored by black. Scale bar: 50 µm

### Rapid freezing preserves inhomogeneous sarcomere patterns under ventricular fibrillation

In addition to the heart showing a regular rhythm, spatiotemporal sarcomere inhomogeneity may be more noticeable in the arrhythmic heart. For example, during ventricular fibrillation (VF), which is characterized by spatiotemporally disordered ventricular excitations and conductions [**29–31**], sarcomere dynamics may be disorganized. As shown in the movies for intracellular Ca^2+^ dynamics in the subepicardial myocardium (**Supplementary Video 2**), while the heart exhibited spatiotemporally uniform Ca^2+^ transients in individual myocytes during regular heartbeats upon excitation (left panel), the VF heart exhibited inhomogeneous Ca^2+^ dynamics, namely, asynchronous, local Ca^2+^ transients with intracellular Ca^2+^ propagating waves within and among individual myocytes, reflecting a chaotic pattern of ventricular excitation (right panel). In accordance with such chaotic Ca^2+^ dynamics, the rapidly-frozen VF heart induced by burst pacing (“RF-VF”, Fig. 7A) showed markedly different SL distributions: individual myocytes extensively exhibited spatially inhomogeneous SLs, showing scattered patterns of a variety of localized SL shortening in accordance with the inhomogeneous Ca^2+^ dynamics observed during VF. In contrast, when the VF heart was fixed with PFA perfusion, the inhomogeneity of SL disappeared (“PFA-VF”, Fig. 7B), similar to the heart with PFA fixation alone (“PFA”, Fig. 6D). As described above, rapid-freezing of the VF heart successfully fixed the heart at the moment of the inhomogeneous contractions/relaxations within and among the individual myocytes reflecting this arrhythmia, whereas such characteristics were entirely lost during the gradual fixation process by PFA perfusion.

**Figure 7.**
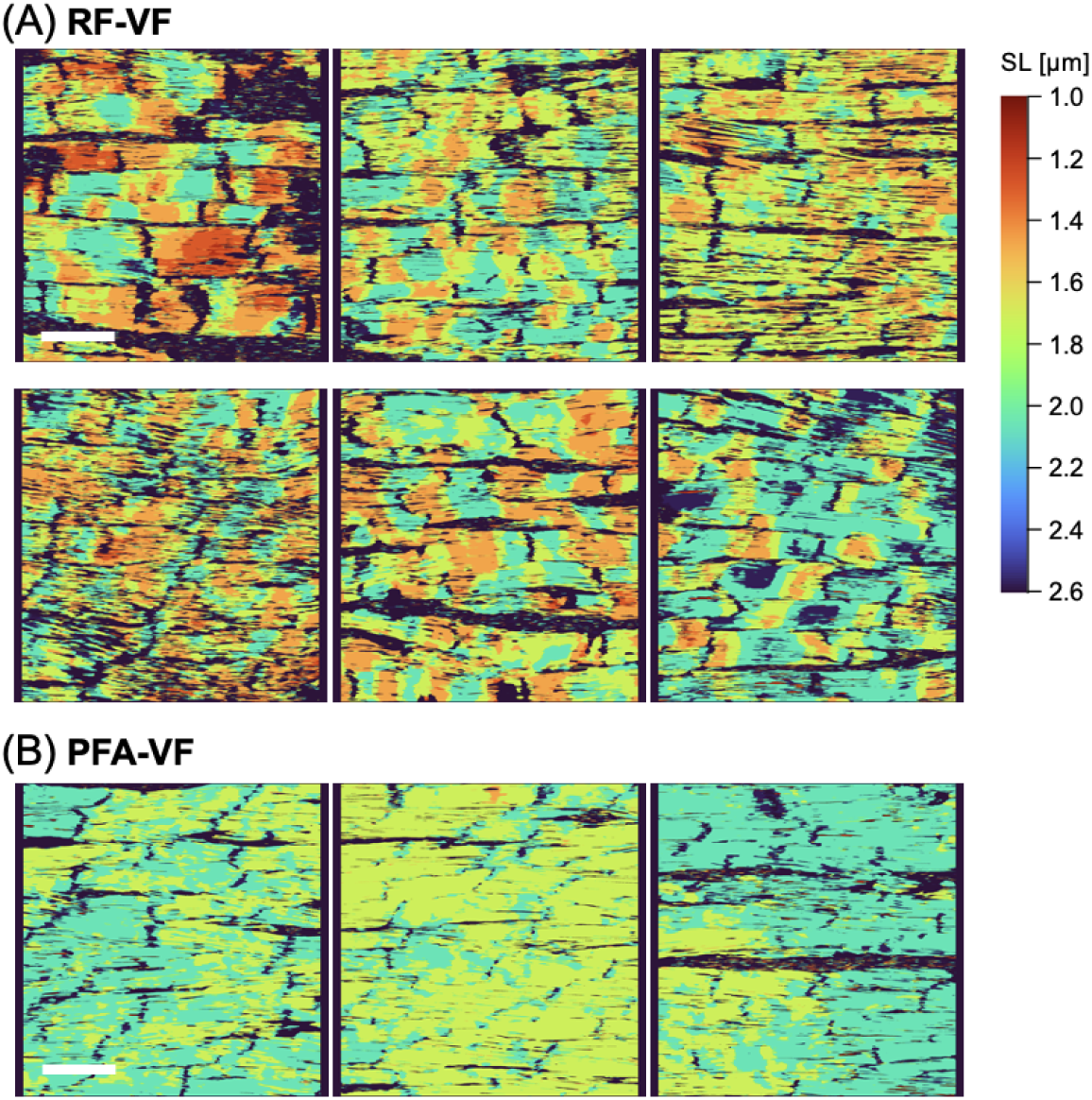
Heatmaps showing the local sarcomere length (SL) in the heart under ventricular fibrillation (VF) fixed with rapid freezing (A: “RF-VF”, n = 2 hearts) and PFA perfusion (B: “PFA-VF”, n = 1 heart). Please note that the areas of nonmyocyte or intra-myocyte SL out of the range of 1.0 to 2.6 µm are colored by black. Scale bar: 50 µm

## Discussion

In this study, we revealed precise cardiac phase-dependent differences in the sarcomere arrangement of myocytes through the rapid-freezing fixation of the Langendorff-perfused rat heart. By adjusting the timepoint of cryogen ejection to the electrically-paced LV surface, we successfully established instantaneous freezing of the heart during peak systole and end diastole. We confirmed that our RF technique and subsequent freeze substitution procedure preserve the cellular and subcellular structures of individual myocytes, especially the sarcomere, which is a fundamental element of mechanical contraction/relaxation [**25**]. Images of immunohistochemical staining for α-actinin in the RF heart revealed marked differences in the SL between systole and diastole: significantly shorter SL was observed during peak systole (mean: 1.57 µm) than during diastole (mean: 1.93 μm). The sarcomeres in the “RF-diastole” heart must be relaxed because their mean SL was nearly similar to that of the “RF-diastole (BDM)” heart (mean: 1.97 µm). Furthermore, we found that the SL value varied within and among the individual hearts even during the diastolic phase. Such variability in SL was attenuated by the pharmacological relaxation of the heart by BDM. Chemical fixation with PFA with or without subsequent RF also decreased the variability of SL, possibly because of the entirely uniform cardiac arrest during gradual fixation by PFA.

For the first time, the variability in the SL was successfully visualized via an SL heatmap of the myocardium, i.e., the phase-specific “snapshot” of the real-time histology for the myocyte sarcomere arrangements in the beating heart, which has not previously been detected by generally used chemical fixation. The SL heatmap for the “RF-systole” heart revealed nearly uniform shortening of SL within and among the individual myocytes, which ranged from 1.4 to 1.8 μm. However, some myocardia showed spatially localized SL shortening, whereas others patchy distributions of varying degrees of SL shortening ranging from 1.8 to 2.1 μm, reflecting spatiotemporally nonuniform myocyte contraction reminiscent of impaired contractile functions. Such spatial inhomogeneity is related to the spatially inhomogeneous myocyte Ca^2+^ transients identified previously under diseased conditions, e.g., metabolic inhibition [**32**] and ischemia [**33**], where individual myocytes exhibit inhomogeneous, localized Ca^2+^ transients with occasional intracellular wave-like propagation as a result of impaired Ca^2+^ release from the sarcoplasmic reticulum [**32, 34**].

On the other hand, the “RF-diastole” heart showing predominant SL elongation also accompanied by patchy distributions of locally short-SL regions, indicating that most of the heart examined during end diastole was not adequately relaxed. In practice, pretreatment of the heart with BDM, which forcibly relaxed the heart [**26**], clearly reduced the number, size, and degree of shorter SL regions, suggesting that the inhibition of actin and myosin interactions by BDM attenuated the localized shortening of sarcomeres. The inhomogeneous SL shortening/elongation has long been identified as a “*tug-of-war*” phenomenon of the sarcomere in the myofibrils [**35, 36**], isolated ventricular myocytes [**37–40**], and individual myocytes directly visualized in the beating heart *in situ* by Kobiruimaki et al. [**41**], who regard this phenomenon as key issues for the harmonization of the myocyte contraction. The present study also directly identified such sarcomere segments through immunohistochemical staining for actin, which showed opposite behaviors from the overall trend, such as the maintenance of gaps even in “systole” or filled gaps in “diastole”.

Spatially inhomogeneous SL shortening may arise in the heart during diastole because a variety of states should be present during diastole, ranging from well-stretched (i.e., highly loaded) conditions to inadequately relaxed (i.e., less loaded) conditions, e.g., diastolic dysfunction. The localized sarcomere contraction during diastole is reminiscent of the occurrence of spontaneous localized Ca^2+^ waves we previously identified as “sporadic Ca^2+^ waves” that arise under quiescent conditions in the perfused rat heart [**7**]. In practice, localized contraction is thought to be caused by the occurrence of spontaneous Ca^2+^ waves, i.e., intracellular wave-like propagations of a localized increase in intracellular Ca^2+^ concentrations in single myocytes, as previously described [**42–44**]. Simultaneous visualization of localized SL shortening with intracellular Ca^2+^ dynamics would confirm this possibility.

We further showed for the first time that the heatmap of the rapidly frozen VF heart exhibits spatially inhomogeneous SL distributions within and among individual myocytes indicating a mixture of localized contractions and relaxations. Such chaotic patterns of SL distributions in the VF heart were diminished by direct chemical fixation of the VF heart; the commonly used chemical fixation procedure is unable to preserve the arrhythmic conditions in the sarcomere arrangement. From the SL heatmaps of rapidly frozen heart and the relevant Ca^2+^ dynamics, the VF heart was found to exhibit inhomogeneous excitation/contraction not only among individual myocytes but also within myocytes.

Despite the clarity of our results, we must consider some of the limitations of this study. First, the present experiments were conducted on an excised, perfused heart with no-blood perfusate, which is quite distinct from a beating heart *in situ*. Second, the two cardiac phases studied were not undergoing physiological beating but underwent low-frequency pacing from the apex. Because of these differences, the spatially variable SL contractions we observed during low-frequency excitations would differ from those in the physiologically beating heart *in situ*; however, we are unable to address this issue because we have no access to a high-performance RF system adequate for high-frequency excitations to more precisely designate the timepoints of the valve opening and the contact of ejected cryogen with the heart surface during high-frequency excitation. In this regard, given that the contact of the ejected cryogen with the heart surface was designed to occur at 30 ms (corresponding to the time point in the 5th CCD frame at 7 ms/frame) after valve opening, the actual time point of cryogen exposure during peak systole ranged from the 5th to 7th CCD frames (i.e., maximum error range of 3 frames or 21 ms) in our experiments. Such inadequate temporal resolution strongly affects the precise timing of cryogen exposure of the heart during high-frequency beating. Our system needs further improvements to solve this issue.

In conclusion, cardiac phase-targeting cryofixation revealed the phase-specific histological behaviors under contractile conditions with spatially variable distributions of sarcomere arrangements in the heart, which have not been identified for beating hearts using current live imaging strategies. Our RF strategy for the heart provides new insights into in-depth spatiotemporal changes in the sarcomere structures for understanding normal and abnormal cardiac functions.

## Methods

### Heart preparation procedures

This study was performed in accordance with the ARRIVE (Animal Research: Reporting of In Vivo Experiments) guidelines 2.0 (https://arriveguidelines.org/). All the animal experiments described in this study were conducted between August 2022 and June 2024 in accordance with *the Guide for the Care and Use of Laboratory Animals* (8th edition, National Academies Press, Washington DC, 2011) following their approval by the Animal Research Committee at Kyoto Prefectural University of Medicine (approval Nos. M2022-238, M2023-231, and M2024-209). Male Wistar rats (9-13 weeks old; n = 25) were purchased from Shimizu Laboratory Supplies (Kyoto, Japan). For the heart excision, after opening the chest of the rat under deep general anesthesia induced by intraperitoneal injections of 0.1 mg/kg medetomidine, 3.0 mg/kg midazolam, and 5.0 mg/kg butorphanol, heparin (1 U/g body weight) was injected into the inferior vena cava, and the heart was then quickly removed and subjected to Langendorff perfusion with HEPES-buffered Tyrode’s solution containing (in mM) NaCl 137, KCl 4, MgCl_2_ 1, NaH_2_PO_4_ 0.33, CaCl_2_ 1.2, HEPES 10 and glucose 10 (pH = 7.4, adjusted by NaOH) for approximately 3 min to wash out the blood. Prior to rapid freezing, the atria were removed to avoid sinus excitation and the atrioventricular junction was mechanically ablated. For rapid cryofixation, the heart was perfused with Krebs-Henseleit (K-H) solution of the following composition (in mM): NaCl 115, KCl 4.0, MgSO_4_ 1.0, NaH_2_PO_4_ 1.2, CaCl_2_ 1.0, NaHCO_3_ 25 and glucose 10 with 95% O_2_ and 5% CO_2_. Perfusion was conducted with a magnetic-driven pump (Micropump 82117, CHUORIKA,.CO.LTD., Mie, Japan) at a flow rate of 5 mL/min. In some experiments 2,3-butanedione monoxime (BDM: 20 mM) (Nacalai Tesque, Kyoto, Japan) was added to the perfusate to attenuate mechanical contraction. For chemical fixation of the heart without rapid freezing or during ventricular fibrillation (VF), the heart was perfused with 2% paraformaldehyde (PFA) for 30 min. The temperature of the perfusate was maintained at room temperature (approximately 23L).

### Rapid-freezing system and protocol

The system, which was designed for the perfused rat heart, was constructed with commercial devices and homemade materials (Fig. 1A, B). Two types of resins (Tough PLA, Ultimaker, Utrecht, the Netherlands and Rigid 10K resin, Formlabs, MA, U.S.A.) and two types of 3D printers (Ultimaker S5, Ultimaker and Form3, Formlabs) were used to fabricate the homemade materials. The main body of the system was composed of two parts: a sample stage (bottom) and a cryogen-ejection system (upper). The stage was composed of a mechanical jack (L490, Thorlabs, NJ, U.S.A.) and a sample box-style chamber to store the cryogen ejected onto the heart (Fig. 1B, top-right panel). The chamber was composed of a slide-glass and spongy blocks as parts of the wall (Fig. 1B, bottom panel) for side-view observation of heart motion and the placement of electrodes, respectively. The perfused rat heart was placed in the chamber with the left ventricular (LV) surface facing upward, where a pedestal fabricated using a 3D printer and parafilm was inserted underneath the heart for cryofixation and subsequent excision of the frozen heart. A pair of silver wire electrodes for electrical stimulation was attached to the apex of the heart. A ring-shaped aluminum electrode was gently mounted on its top plane for electrocardiogram (ECG) recordings and detection of the time at which the cryogen reached the heart.

The cryogen-ejection system was composed of a tank, a barometer (B15170, Nisshin Gauge MFG. CO., Ltd, Osaka, Japan), and an electromagnetic valve (B2067-LN2-LB-5VDC, Fortive ICG Japan Co., Ltd., Osaka, Japan), which were assembled into a block made with a 3D printer. The ejection system was mounted on the sample stage using a small mechanical jack (L200, Thorlabs) to adjust the distance (approx. 4 cm) between the ejection system and sample stage. Heart contraction was evoked by electrical stimulation at 0.5 Hz with a pulse width of 1 ms and amplitudes of 20-30 mA (corresponding to the suprathreshold current) applied to the apex through a stimulator (ESTM-9; Brainvision, Tokyo, Japan) and an isolator (A385, World Precision Instruments, FL, U.S.A.), which were regulated by PC software (BV ESTM-9, Brainvision). The perfusate that leaked from the heart was continually collected with a suction tube placed in the chamber using a peristaltic pump (MP-2000, TOKYO RIKAKIKAI Co., Ltd., Tokyo, Japan). A side-view movie of heart contraction was recorded with a charge-coupled device (CCD) camera (HR, Brainvision: sensor size of 6.4 mm × 4.8 mm, pixel number of 375 × 252 pixels, frame rate of 143 frames/s) with a camera lens (LM25JC5M2, Kowa Optronics, Aichi, Japan) through the glass wall of the chamber. The ECG of the heart was acquired with a reference electrode and a ring-shaped electrode using amplifier (UA-200 loaded in the power-supply unit USB-100, Unique Medical, Tokyo, Japan) and a low-pass filter (LPF-100A: Warner Instruments, MA, U.S.A.) at sampling rate of 2857 points/s.

Liquid propane was poured into the tank as a cryogen in the cryogen-ejection system, which was pressurized to 0.05 MPa above atmospheric pressure by N_2_ gas injection while the cold temperature was maintained with liquid nitrogen. Under recording of the side-view movie and ECG, the heart was rapidly frozen by ejecting the cryogen after the electromagnetic valve was opened at the preset time-point (Fig. 1C). Valve opening was regulated by a TTL trigger signal output from a stimulator (ESTM-9, Brainvision) and amplified by an electrical supplier (LEDD1B, Thorlabs). The rapidly-frozen heart was stored in the deep freezer (−80L) until the freeze substitution procedure. For a detailed protocol in the rapid-freezing process, see **Supplementary Methods** online.

The side-view images of the heart and ECG with electrical stimulation signals were simultaneously collected to an image acquisition system (MiCAM02, Brainvision). The saved data were analyzed using BV Ana (Brainvision) and the open-source software Fiji (National Institute of Health, MD, U.S.A.) [**45**]. The aspect ratio of the CCD images was compensated using the “HR” function of BV Ana. Sequential changes in heart contraction were estimated by as the vertical motion of the ring-shaped electrode.

### Freeze substitution

Freeze substitution was performed using a previously reported method [**21**] with minor modifications. The freeze substitution solution was made by adding 20% paraformaldehyde (PFA: 16005-1KG-R; Sigma-Aldrich/Merck KGaA; Darmstadt, Germany) in distilled water to pure acetone (013-00356; FUJIFILM Wako Pure Chemical Corporation, Osaka, Japan) at a final PFA concentration of 2%. The solution was completely dehydrated using a molecular sieve (3A, 134-06095; FUJIFILM Wako Pure Chemical Corporation), and then stored in a deep freezer (−80 L). The rapidly-frozen heart in the deep freezer was immersed in the chilled freeze-substitution solution for 48 hours in deep freezer.

Thereafter, temperature of the heart in solution was gradually elevated from −30 L, −10 L, and 4 L at 2-hour intervals to room temperature at 23 L. During this process, the heart was gradually fixed with PFA and acetone while the structures fixed by rapid freezing was maintained. The freeze-substituted heart was subsequently washed with pure acetone 3 times for 30 min each, and the rapidly frozen region (a part of the left ventricular surface located inside the ring-shaped electrode) was gently cut from the heart using surgical knives at room temperature. Finally, the tissue blocks were immersed in phosphate-buffered saline (PBS) for HE- or immunofluorescence staining.

### Preparation and observation of HE-stained thin section

Heart tissue blocks were paraffinized using a standard protocol with ethanol and xylene. The paraffinized tissue blocks were sliced into 4-µm thick sections with a microtome to generate cross-sections of the rapidly frozen and nonrapidly frozen region or sections in the myocardial planes near the heart surface (within 30 µm) underneath the rapidly frozen region. After the sections were attached to glass slides and deparaffinized using a standard protocol, they were stained with hematoxylin and eosin. Bright-field images of the HE-stained thin sections were recorded using a digital slide scanner (NanoZoomer S360, Hamamatsu Photonics, Hamamatsu, Japan), and analyzed on NDP.view2 software (Hamamatsu Photonics) and Fiji (National Institute of Health).

### Preparations for immunohistochemistry

The tissue blocks were immersed in 0.1% Triton X-100 (28314, Thermo Fisher Scientific, IL, U.S.A.) in PBS for 30 min at room temperature. After three washes with freshly prepared PBS, the blocks were immersed in 10% skim milk (198-10605; FUJIFILM Wako Pure Chemical Corporation) in PBS for 30 min at room temperature and then washed with freshly prepared PBS three times. For-fluorescence staining, the sections were subsequently incubated with combinations of primary antibodies (24−48 hours at 4L) and secondary antibodies (24−36 hours at RT, after three washes with freshly prepared PBS), as described below.

For visualization of subcellular structure of the myocytes, an anti-α-actinin antibody (1:400, mouse monoclonal IgG, A7811, Sigma-Aldrich/Merck) and anti-titin antibody (1:400, rabbit polyclonal IgG, 27867-1-AP, Proteintech, IL, U.S.A.) were used as primary antibodies to stain the Z-line and M-line in the sarcomeres of myocytes. An antibody against cardiac myosin heavy chain (1:400, mouse monoclonal IgG [BA-G5], ab50967, Abcam, Cambridge, United Kingdom) was used to stain the thick filaments. As secondary antibodies, IgG H&L Alexa Fluor 555 (1:300; ab150114, Abcam) was used to label the Z-line and thick filaments, and goat anti-rabbit IgG H&L Alexa Fluor 488 (1:300, ab150077, Abcam) was used to stain the M-line. The antibodies were diluted in PBS supplemented with 1% bovine serum albumin (BSA, 017-15141, FUJIFILM Wako Pure Chemical Corporation) and 0.1% Tween 20 (28230, Thermo Fisher Scientific). Additionally, thin filaments (F-actin) and cell membranes were labeled with CytoPainter Phalloidin-iFluor 405 Reagent (1:500 in PBS; ab176752, Abcam) and WGA Alexa Fluor 633 conjugate (1:500 in PBS, W21404, Thermo Fisher Scientific), respectively, immune-fluorescence staining process and three washes with freshly prepared PBS.

For quantification of the SL, an anti-α-actinin antibody (1:500 in PBS) was used as a primary antibody to stain the Z-line, and goat anti-mouse IgG H&L Alexa Fluor 488 (1:250 in PBS; A-11001, Themo Fisher Scientific) was used as a secondary antibody. For some heart specimens, the intercalated discs between myocytes were also stained with an anti-N-cadherin antibody (1:200 in PBS, rabbit polyclonal IgG, ab76011, Abcam) and goat anti-rabbit IgG H&L Alexa Fluor 555 (1:250 in PBS, A-21422, Thermo Fisher Scientific). After three washes with freshly prepared PBS, the cell membranes were labeled with a WGA Alexa Fluor 633 conjugate (1:100 in PBS; W21404, Thermo Fisher Scientific).

The stained heart blocks were washed with freshly prepared PBS three times, and then placed in glass-bottom dishes (D11140H, Matsunami Glass Ind., Ltd., Osaka Japan) with anti-bleaching mounting media (H-1900, Vector Laboratories, CA, U.S.A.).

### Fluorescence imaging with confocal laser-scanning microscopy

Fluorescence imaging of the heart block was conducted using confocal laser-scanning fluorescence microscopes. For fluorescence imaging of the subcellular structures of the myocyte, an LSM 900 (Carl Zeiss, Oberkochen, Germany) was used in confocal imaging mode. An oil-immersion objective lens (63×/1.40, Plan-Apochromat, DIC M27, Carl Zeiss) was used. The excitation wavelengths for each target were 405 nm for thin filaments (F-actin, CytoPainter Phalloidin-iFluor 405 Reagent), 488 nm for the M-line (titin, AlexaFluor488), 561 nm for the Z-line (α-actinin, AlxaFluor555) and thick-filaments (myosin heavy chain, AlexaFluor555), and 640 nm for the cell membrane (WGA-AlexaFluor633 conjugate). The obtained image size ranged from 112.68 µm × 112.68 µm (1596 × 1596 pixels) to 126.77 µm × 126.77 µm (1796 × 1796 pixels), with a fixed pixel size of 0.071 µm × 0.071 µm. The pixel dwell time ranged from 9.40 to 10.58 µs, and the pinhole size was 50 µm, corresponding to 1.00-1.26 Airy unit size depending on the excitation/detection channel, gain and offset of the detector were 500 V and 0, respectively, and the digital gain was 1.0. For the imaging of α-actinin immunofluorescence staining to quantitatively analyze the sarcomere length, an FV1000 confocal laser scanning microscope (Olympus/Evident, Tokyo, Japan) was used. An oil-immersion objective lens (60×/1.42, PLAPON, Olympus/Evident) was used, and the excitation wavelength for each target was 488 nm for the Z-line (α-actinin, Alexa Fluor 488), 543 nm for intercalated disk (N-cadherin, Alexa Fluor 555), 633 nm for cell membrane (WGA-Alexa Fluor 633 conjugate). The obtained image size was 210.84 µm × 210.84 µm (2048 × 2048 pixels), with a pixel size of 0.103 µm × 0.103 µm. The pixel dwell time was 20.0 µs, the pinhole size was 150 µm, and the detector gain was 500 – 520 V.

All of the obtained fluorescence images were analyzed (cropping, merging of color channels, and line profiling) with Fiji (National Institute of Health). The detected line-profiles were analyzed using Igor Pro 9 (WaveMetrics, OR, U.S.A.).

### Quantification of the SL

The sarcomere length of each myocyte was calculated by discrete Fourier transform (DFT) to determine the periodical pattern of striations in each cell, which was obtained from the intensity line profiles along the longitudinal axis of a cell on the images of α-actinin immunofluorescence staining and a cellular contour line (WGA and/or N-cadherin staining). The process was based on a previously reported method [**9**] with some minor modifications to fit our experimental conditions, and was performed using a home-built software constructed with MATLAB (R2024a, MathWorks, MA, U.S.A.). A line was drawn on each cardiomyocyte by a user, approximately perpendicular to the periodical pattern of striations within the cellular contour in images of α-actinin immunofluorescence staining. The length of the line was regulated such that approximately more than two-thirds of the maximum width (along perpendicular to the striations) of each targeted cell was filled and fit in the range of 512 to 1024 pixels (corresponding to 52.7-105.5 µm on the image scale). In this step, cardiomyocytes corresponding to either of the following conditions were excluded from the target: (i) those whose contour line or cellular body was disrupted by the edge of field of view or by the overlaps of extracellular structures, and (ii) those whose maximum width was shorter than 512 pixels. Next, the user assigned a travel distance for a vertical shift of the drawn line to fill approximately more than half of the cellular height. Afterwards, the drawn line was vertically shifted by a pixel from top to bottom along the assigned travel distance, and at each location, the line was rotated by 1° within ± 5°. Through the vertical shift and rotation of the line, the intensity-profile was detected as a mean of intensity-profiles along the line with ±1 pixel in the vertical direction, at each location and rotated angle (with a compensation using the nearest neighborhood approach approach). For each detected intensity-profile, an amplitude spectrum was calculated using a “fft” function based on the DFT of MATLAB with zero-padding to 1024 points. The horizontal axis for the amplitude spectrum was converted to cycle length (i.e. sarcomere length) and was cropped to the range of 1.1 to 2.7 µm. A peak ratio between the amplitude of the highest peak and the second highest peak on the cropped range was calculated, and tentatively recorded along with the peak position and amplitude of the highest peak. After the calculation for all locations and rotated angles of the line, an amplitude spectrum with the peak of the highest amplitude was selected among the spectra satisfying a peak ratio greater than 3, and the peak position on the cycle length (µm) was assigned to the targeted cell as its representative SL; the corresponding line location and rotated angle, intensity-profile, and amplitude spectrum were recorded. If no amplitude spectrum satisfying a peak ratio greater than 3 was observed for a cell, the cell was excluded from the analysis. The representative intensity profiles and amplitude spectra in this paper were displayed after the normalization of the differences between the highest and lowest values.

### Statistics

The statistical significance of differences in the SL per heart was estimated by using 1-way analysis of variance (ANOVA) followed by Tukey post-hoc test for four-group comparison (Fig. 5B) and student t-test for two-group comparison (**Supplementary** Fig. 4A) (Prism, ver. 10.1.2, GraphPad Software, MA, U.S.A.). For the analysis of the SL per cell, the linear mixed effects (LME) model was applied to the database to reconstruct the hierarchical data set with fixation conditions as a fixed effect and the individual heart as a random effect, thus avoiding pseudoduplication by data points collected from the same heart [**27, 28**]. The statistical significance of differences in sarcomere length per cell was subsequently estimated by ANOVA followed by the Tukey test for four-group comparison (Fig. 5C) and t-test based on the Satterthwaite approximation for two-group comparison (**Supplementary** Fig. 4B). The application of the LME model and subsequent ANOVA/Tukey test were conducted using R (ver. 4.5.0, R Foundation for Statistical Computing, Vienna, Austria) coded in RStudio (ver. 2025.05.0+496, Posit PBC, MA, U.S.A.). All the graphs for the statistical results shown in this paper were prepared using Prism software. Differences with a probability of 5% or less were considered significant (PL<L0.05).

### Heatmap reconstruction

Using the images of α-actinin immunofluorescence staining, heatmaps were generated to visualize the local distribution of the SL within and among the cardiomyocytes in the cryofixed hearts, with a custom-built software coded with MATLAB (R2024a, MathWorks). The heatmap was generated by replacing the fluorescence intensity values for each pixel in the fluorescence image with the values of SL determined using surrounding periodical information, and the image was converted to a color map of the SL values. The SL value assigned to a pixel ((*x,y*) in the fluorescence image) was determined from an amplitude spectrum calculated by DFT (“fft” function in MATLAB) for an averaged intensity profile along the horizontal direction of the local region with a width of 100 pixels and a height of 3 pixels (*x - 50*: *x + 49, y-1*: *y +1*). In the amplitude spectrum for the cycle length axis of the myocytes (µm), the position of the maximal amplitude was adopted as a representative SL. If the calculated representative-SL value was out of the range of 1.0 to 2.6 µm, 0 was assigned to the corresponding pixel instead of the SL value. DC removal and a window function (symmetric Hann window) were applied to the profile in advance of the DFT processing to suppress the edge effect of the averaged intensity profile to the frequency domain. The number of pixels in the fluorescence image was (*x*) 2048 × (*y*) 2048 pixels, and the SL value (i.e. the DFT processing) was assigned to the center (*x*) 1948 × (*y*) 2046 pixels in the image by taking margins for the local region at the edge of the image. After the pixel values were converted, all the values were inverted and normalized, and a color table of “turbo” was applied so that the smaller SL values (shorter SL) were indicated as hotter colors on the map.

### Rapid-scanning confocal Ca^2+^ imaging

Ca^2+^ imaging of the perfused rat heart was performed essentially as described in our previous studies [**7, 46**]. The excised heart was subjected to Langendorff perfusion with HEPES-buffered Tyrode’s solution and subsequently incubated with the Ca^2+^ indicator, fluo4-AM (5.5 μM, Dojindo, Kumamoto, Japan) for 15 min at room temperature (23-25 L). After fluo4 loading, the heart was perfused with HEPES-buffered Tyrode’s solution containing 1 mM Ca^2+^ and probenecid (0.1 mg/ml) at 37 L for 5 min for de-esterification of the acetoxy methyl ester (AM) form of the fluo4. The fluo4-loaded heart was then used in subsequent experiments and maintained under constant perfusion with 0.3-mM Ca^2+^-containing K-H solution (5 mL/min) at 30 L using a tube heater (Kawai Corporation, Aichi, Japan). ECGs of the hearts were recorded using silver wires (0.5 mm diameter) placed at the bottom of a customized heart chamber. The hearts were treated with 15 µM of blebbistatin prior to measurement to suppress the oscillation of field of view by the heart during imaging.

We conducted rapid-scanning confocal imaging of the fluo4 fluorescence intensity (251 × 196 μm, 376 × 252 pixels, 100 frames/s) in the subepicardial myocardium of the LV using constant Langendorff perfusion with 0.3 mM Ca^2+^ K-H solution. The images were captured with a high-speed, multipoint-scanning confocal microscope constructed from an upright microscope (BX-50WI, Olympus/Evident) and a spinning disc-type confocal unit CSU-21 (Yokogawa, Tokyo, Japan). The fluo4-fluorescence signals were amplified using an image intensifier (C8600, Hamamatsu Photonics, Shizuoka, Japan) and captured using a CCD camera (HR, Brainvision) with a detection system (MiCAM02, Brainvision). The images were captured using a 20× objective lens (UMPLan FI, NA = 0.5, Olympus/Evident), and some experiments included electrical stimulation of the left ventricular apex using silver wires (0.5 mm in diameter). The obtained images were postprocessed using BV Ana (Brainvision) and Fiji (National Institute of Health). The aspect ratio of the images was compensated by “HR” function of BV Ana. After the intensity in all the images was normalized by subtracting the intensity in a reference image consisting of minimum intensity at each pixel through the measurement, the images were cropped to 330 × 250 pixels (220 × 167 µm^2^).

### Induction of ventricular fibrillation (VF)

VF was induced by burst pacing (at 20 Hz for 120 s) to the heart from the ventricular apex under Langendorff perfusion with low K^+^ (1 mM) K-H solution [**47**]. After 3 min of observation to confirm that the VF was stable, rapid freezing of the VF heart was conducted by rapid exposure to the cryogen using our ejecting system without determining the time point of cryogen exposure.

## Data availability

All the data in this paper are available from the corresponding authors upon request.

## Acknowledgements

This work was partially supported by the Core Research for Evolutionary Science and Technology (CREST) fund (JPMJCR1925) from the Japan Science and Technology Agency (JST), and Grant-in-Aid for Early-Career Scientists (No. 22K18174) of the Japan Society for the Promotion of Science.

## Author contributions statement

S.T. and K.M. performed the experiments and analyzed the data, and wrote the first draft of the manuscript. Y.K., M.Y. and K.F. designed and fabricated the main body of the rapid-freezing system. S.T., K.M., Y.M., Y.H., and H.T. contributed to the construction of the integrated system for phase-targeting rapid-freezing. Y.M. and W.J.H. supplied a part of supplementary data. K.M. and H.T. supervised the work, and H.T. contributed to the final manuscript approval. All the authors reviewed and approved the final manuscript.

## Additional information

The authors declare no competing interests.

