## Supplementary Information for "Phase-targeting rapid cryofixation of the beating heart and histological analysis unveil contractile state-dependent sarcomere dynamics"

**Supplementary Figures 1 to 5**

**Legends for Supplementary Movies**

**Supplementary Methods**

### Supplementary figures

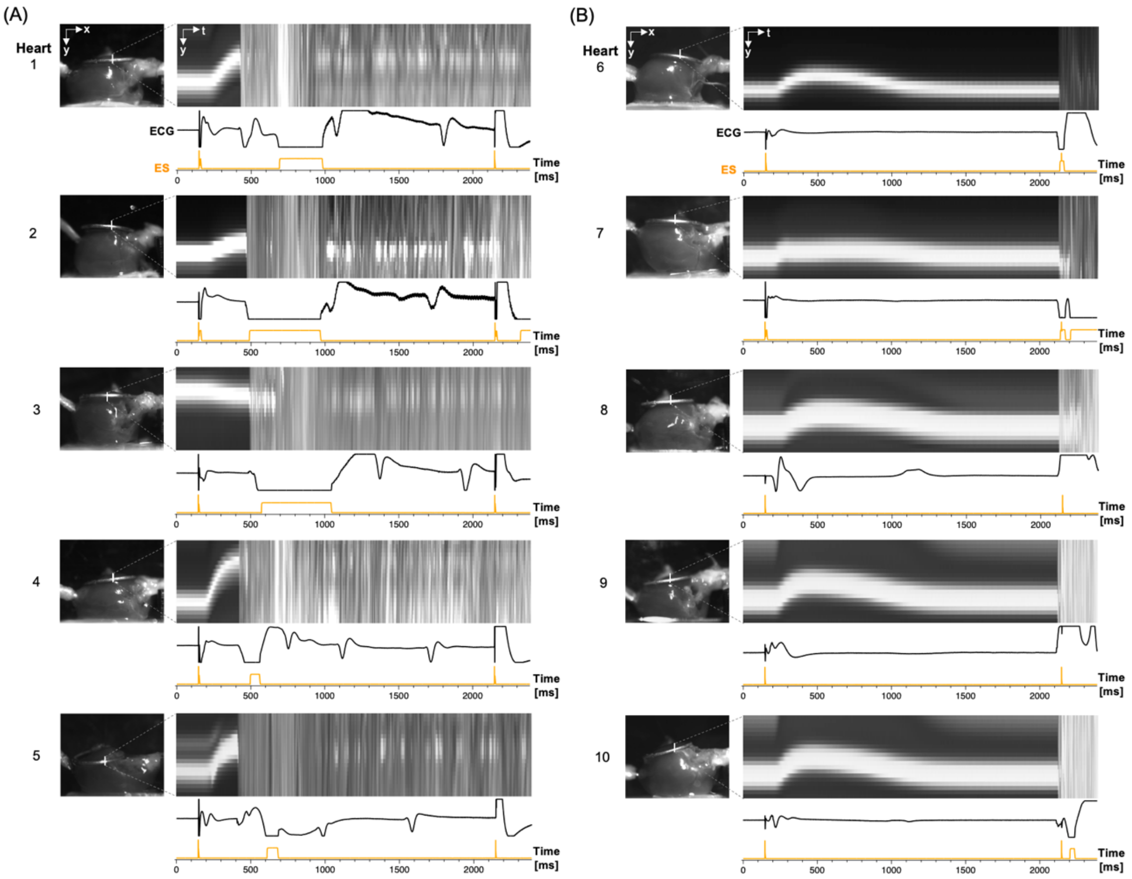


**Supplementary Figure 1.** Side-view images and vertical-shift images (sequential traces for the motions of the ventricular surface (ring electrode)) with rapid-freezing during (A) peak systole and (B) end diastole with electrocardiogram (ECG) and electrical stimulation (ES, 0.5 Hz). Please note that the heart 3 exhibited downward-moving of the ventricular surface during systole while the other ones showed upward-moving.

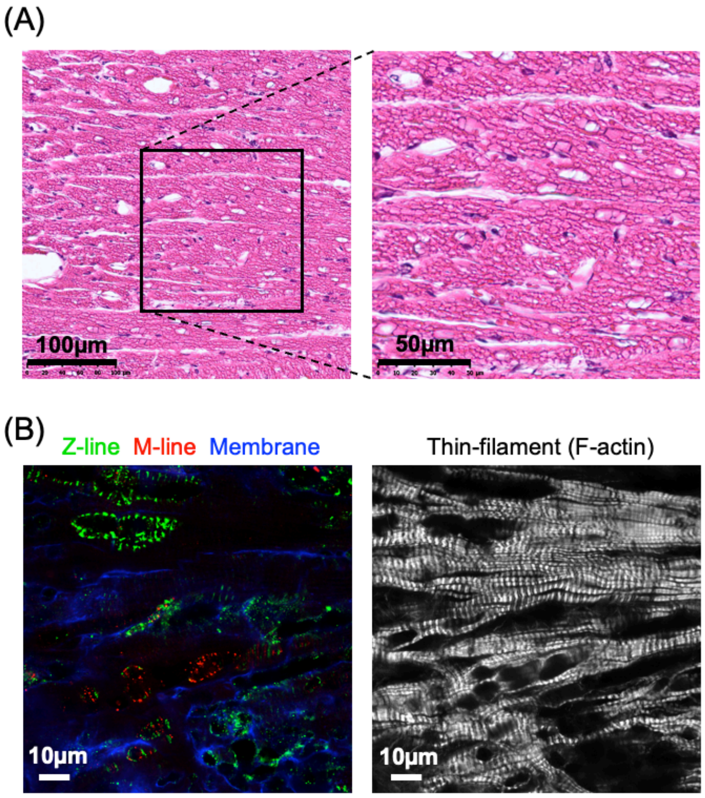


**Supplementary Figure 2.** Bright-field images of the HE-stained thin sections (A) and fluorescence images of an immunostained block (B) of the heart, which was once thawed at room temperature during the rapid-freezing process. The thickness of the HE-stained section is 4 μm. Fluorescence images were obtained with a confocal laser-scanning fluorescence microscope. The distributions of the Z-lines, M-lines, membranes, and thin filaments were obtained as results of immunostaining for α-actinin/titin and staining with WGA/phalloidin, respectively. The vacuoles observed in both the HE-stained and fluorescence images are probably caused by ice-crystal formation during the thawing and refreezing processes.

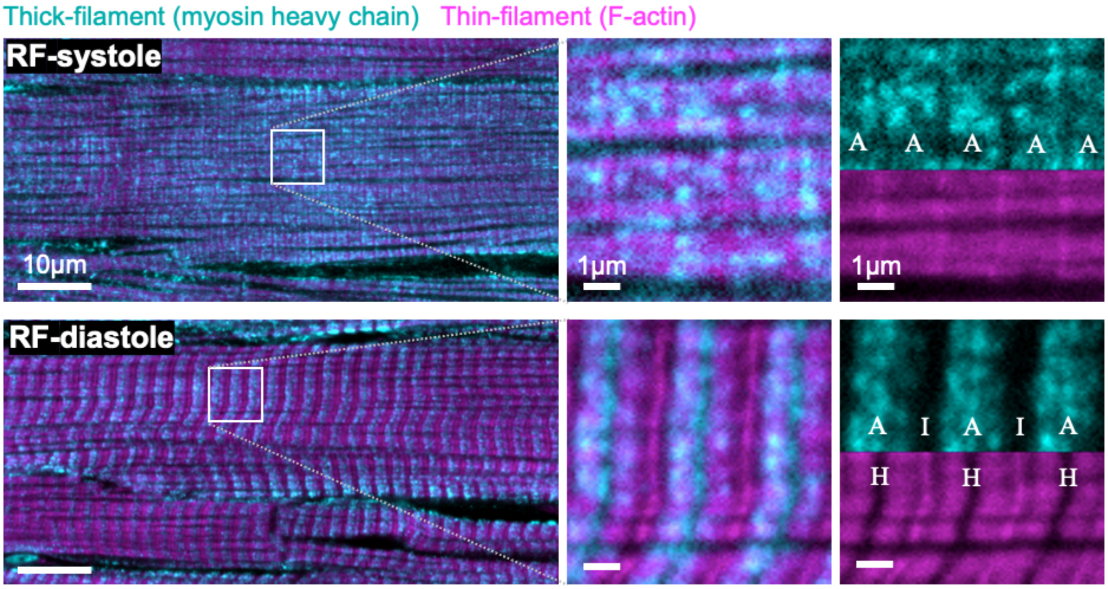


**Supplementary Figure 3.** Fluorescence images of thick-filaments (myosin heavy chain) and thin-filaments (F-actin) labeled by immunofluorescence staining for myosin and fluorescence staining for phalloidin, respectively. The images in the middle are enlarged from the regions in the white squares in the images on the left. The right panels shown color-separated images of the corresponding enlarged region. The positional relation of thick- and thin-filaments reveals the location of the A-band (corresponding to thick filaments), I-band (the region of thin-filaments not overlapping with thick filaments) and H-band (the region of thick-filaments not overlapping with thin filaments), especially for "RF-diastole" hearts.

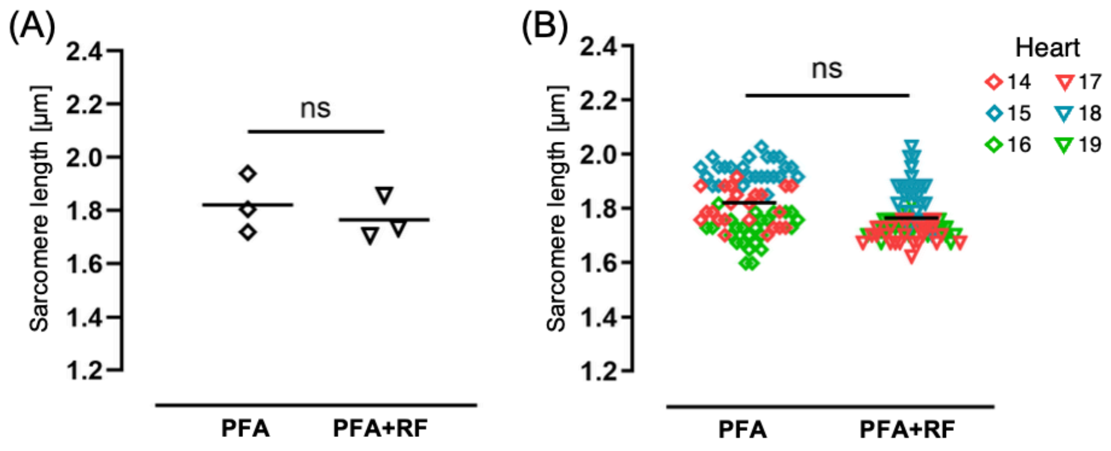


**Supplementary Figure 4**. Results of statistical analysis for the sarcomere length of the heart fixed with PFA-perfusion ("PFA") and rapid-freezing after PFA fixation ("PFA+RF"). Sarcomere lengths per heart (A: n = 3 hearts) and per myocyte (B: n = 20-25 cells/heart).

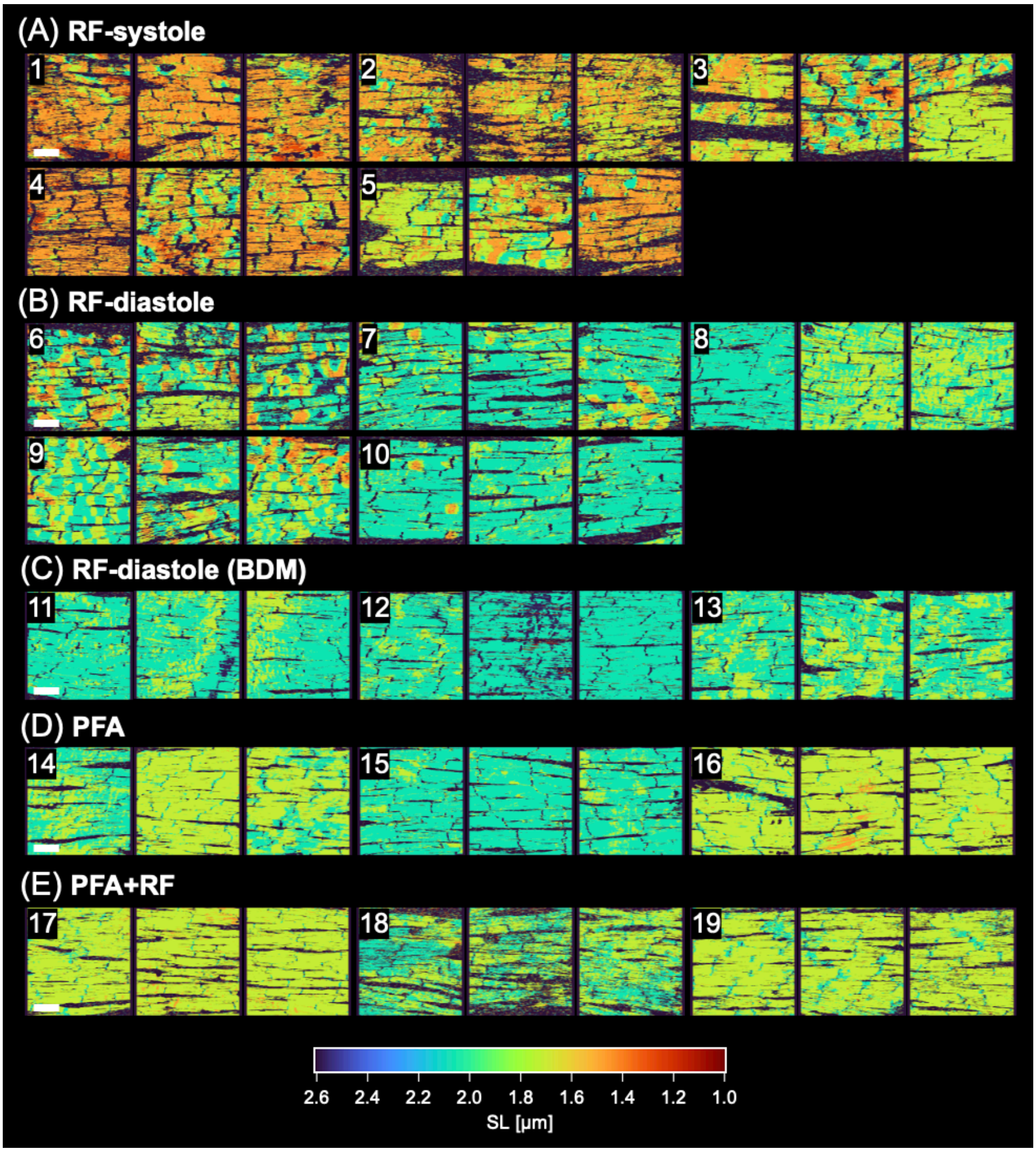


**Supplementary Figure 5**. Heatmaps of all the hearts. (A) Rapid freezing during peak systole ("RF-systole"), (B) rapid freezing during end diastole ("RF-diastole"), (C) rapid freezing during end-diastole under BDM treatment ("RF-diastole (BDM)"), (D) fixation with paraformaldehyde (PFA) perfusion alone ("PFA"), and (E) rapid freezing after fixation by PFA perfusion ("PFA+RF"). The numbers in the images indicate the IDs of the hearts. Scale bar: 50 μm.

### Legends for Supplementary Videos

**Supplementary Video 1.** Movies of the motion of the heart in side view upon rapid freezing during peak systole ("RF-systole") shown in Fig 2A. Frame rate: 143 frames/s (7.0 ms/frame) for the recording and 50 frames/s for the display.

**Supplementary Video 2.** Movies of $Ca^{2+}$ fluorescence imaging of subepicardial myocardium and ECG of perfused rat hearts (A) during regular heartbeat and (B) during ventricular fibrillation (VF). The displayed fluorescence images were prepared by subtracting the intensity from reference images constructed from the minimum intensity at each pixel through imaging. Frame rate of fluorescence imaging: 100 frames/s for the recording and 50 frames/s for the display.

### Supplementary Methods

#### Protocol for the phase-targeting rapid-freezing of the Langendorff-perfused rat heart

In the working space for this experiment, adequate ventilation and oxygen/propane monitoring were ensured to prevent the accumulation of evaporated propane refrigerant or liquid nitrogen. We prepared approximately 50 mL propane as a cryogen, which was liquefied and maintained at a low temperature (approximately -190 °C) in a glass beaker immersed in liquid nitrogen until use, for the rapid-freezing of a heart. The heart, which was placed in a sample chamber of the rapid-freezing system with the left ventricle facing upward, was electrically stimulated at 0.5 Hz with a pair of silver electrodes inserted into the ventricular apex. The ECG was recorded from the electrodes: one from the ring-shaped electrode placed on the upper surface of the heart and another on the bottom.

Liquid nitrogen was poured into the compartment for the electromagnetic valve in the cryogen ejection system to cool the valve (*see* Fig. 1A). The timing for cryogen ejection was determined from side-view video images of heart motion and ECG with electrical stimulation (ES) signals recorded in advance. For the rapid freezing of the heart during peak systole, a time lag [α] ms from an electrical stimulation to peak contraction of the heart was estimated. In our experiments, the [α] ranged between 250 to 350 ms by the five different hearts. By considering another time lag from a TTL-trigger output for valve opening to the contact of ejected cryogen and the heart, which was preliminarily estimated as 30 ms, the TTL trigger in the rapid-freezing process was initially set to fire at the time point of [α-30] ms from an electrical stimulation after starting electrical stimulation and recording. Additionally, we added a margin of around 10 ms to α to avoid contact between the ejected cryogen and the heart surface before its peak systole which lasted several hundred milliseconds. For the rapid freezing of the heart during end diastole, the TTL trigger for valve opening was set to fire at the time point of 1920 ms (30 ms prior to the target time point of 1950 ms) from an electrical stimulation. After setting the time point for the TTL trigger output and just before starting rapid-freezing process, the tank in the cryogen ejection system was filled with prepared cryogen (approximately 25 mL) sealed with a gauge pressure of 0.05 MPa by injecting nitrogen gas. After that, the rapid-freezing process by the ejection of cryogen was performed by starting the electrical stimulation and recording of side-view images and ECG. All the procedures from the filling of liquid nitrogen in the compartment to the ejection of cryogen were completed within 5 minutes so as not to hinder the valve opening by supercooling.

After the rapid-freezing process, the ejected cryogen filled up the heart entirely in the sample chamber. As needed, the cryogen was additionally poured into the chamber manually to avoid exposure of the frozen heart to the air. After turning off the nitrogen-gas injection to the tank, the frozen heart was quickly transferred in the liquid nitrogen. In this transferring process, the ring-shaped electrode was removed, and the perfusion cannula and stimulation electrode, both of which were inserted into the frozen heart, were cut out. Finally, the frozen heart was transferred into a 50 mL conical tube, which was kept cool by liquid nitrogen, the stored in a deep freezer (-80℃) until the freeze substitution procedure.
